# Epigenetic Landscape of HIV Infection in Primary Human Macrophage

**DOI:** 10.1101/2022.01.25.477798

**Authors:** Fang Lu, Yanjie Yi, Olga Vladimirova, Urvi Zhankharia, Ronald G. Collman, Paul M. Lieberman

**Affiliations:** The Wistar Institute, Philadelphia, PA 19104; University of Pennsylvania Perelman School of Medicine, Philadelphia, PA

**Keywords:** HIV, latency, macrophage, microglia, histone, chromatin, DNA methylation, hydroxymethylation

## Abstract

HIV-infected macrophages are long-lived cells that sustain persistent virus expression, which is both a barrier to viral eradication and contributor to neurological complications in patients despite antiretroviral therapy (ART). To better understand the regulation of HIV in macrophages, we compared HIV infected primary human monocyte derived macrophages (MDM) to acutely infected primary CD4 T cells and Jurkat cells latently infected with HIV (JLAT 8.4). HIV genomes in MDM were actively transcribed despite enrichment with heterochromatin-associated H3K9me3 across the complete HIV genome in combination with elevated activation marks of H3K9ac and H3K27ac at the LTR. Macrophage patterns contrasted with JLAT cells, which showed conventional bivalent H3K4me3/H3K27me3, and acutely infected CD4 T cells, which showed an intermediate epigenotype. 5‘-methylcytosine (5mC) was enriched across the HIV genome in latently infected JLAT cells, while 5‘-hydroxymethylcytosine (5hmc) was enriched in CD4 and MDM. HIV infection induced multinucleation of MDMs along with DNA damage associate p53 phosphorylation, as well as loss of TET2 and the nuclear redistribution of 5-hydoxymethylation. Taken together, our findings suggest that HIV induces a unique macrophage nuclear and transcriptional profile, and viral genomes are maintained in a non-canonical bivalent epigenetic state.

**Importance:** Macrophages serve as a reservoir for long-term persistence and chronic production of HIV. We found an atypical epigenetic control of HIV in macrophages marked by heterochromatic H3K9me3 despite active viral transcription. HIV infection induced changes in macrophage nuclear morphology and epigenetic regulatory factors. These findings may identify new mechanisms to control chronic HIV expression in infected macrophage.

## Introduction

HIV results in lifelong infection requiring continuous antiretroviral therapy (ART) to suppress viral replication and prevent immune deficiency (1). A major barrier to cure is the existence of long-lived infected cells that persist during ART. Multiple anatomic sites may contribute to viral persistence including blood, lymphoid tissue, gut-associated lymphoid tissue, bone marrow, and brain (2–4). While most attention has focused on CD4+ T cell reservoirs, myeloid cells are well-established to harbor virus in the CNS, where macrophages and microglia are the principal infected cell type (5–7). Functional cure therefore requires attention not only to CD4+ T cell reservoirs but to other cells, including myeloid reservoirs.

Monocytes are generally resistant to HIV infection but become permissive as they mature into macrophages (8, 9). Macrophages are thus terminally differentiated non-dividing cells yet susceptible to robust HIV infection (10, 11). This feature differentiates them from T cells, which require activation and cell proliferation to become robustly susceptible in vitro (although limited low-level infection of resting T cells may be achieved in some models (12–14)). Thus, macrophage infection and establishment of integrated provirus occurs in the context of a unique cellular microenvironment relative to T cells, particularly with regards to chromatin structure. Another distinguishing feature of macrophage infection is that they are long-lived cells yet resistant to HIV-induced killing, and productively infected macrophages persist for prolonged periods, unlike activated CD4^+^ T cells that are killed by active virus replication (15–17). This feature enables infected myeloid cells to serve as long-term reservoirs in vivo, particularly in the CNS (18–21). Finally, while the resting CD4^+^ T cell long-term reservoir is typically thought of as latent, low-level virus expression generally persists throughout the lifespan of infected macrophages (18, 22–24). Indeed persistent low level virus expression from long-lived infected brain macrophages is thought to be a driver of neurological complications that occur in infected people despite ART (25, 26). Thus, macrophages likely control HIV differently than CD4^+^ T cells and studies of epigenetic control in T cell models may not be sufficient to understand persistent infection in differentiated primary macrophages.

Transcriptional and epigenetic regulation of HIV in T-cells has been studied extensively (27, 28). Transcriptional regulation occurs primarily at the 5’ LTR and is mediated by various transcription factors that control the modification and positioning of nucleosomes that restrict transcription initiation and elongation (27, 29, 30). In contrast to T cells, less is known about epigenetic factors that control HIV infection in macrophages. Here, we investigate epigenetic features, including active and repressive histone marks and DNA methylation and hydroxymethylation status across the HIV genome in infected primary human monocyte-derived macrophages (MDMs) that regulate HIV genome in post-integration latency. We compare MDMs to productively infected primary CD4^+^ T cells, and also with an established latency model in Jurkat T cells using JLAT cells. Our results revealed surprising non-canonical bivalent chromatin structures of HIV genome in primary infected macrophages.

## Results

### Epigenetic profiles of HIV genomes in primary macrophages, primary CD4 T cells and latent cell line JLAT

The epigenetic modification of histones associated with HIV genomes in MDM, CD4 T cells, or latently infected T-cell line JLAT 8.4 was investigated by ChIP-qPCR assay using primers sampling regions across the entire viral genome (**Fig. 1**). Primary MDM and CD4 T cells were infected with the brain-derived macrophage-tropic HIV-1 primary isolate YU2 (31), while JLAT 8.4 cells carry a single clonally integrated HIV genome derived from a prototype strain (32). We assayed for histone H3K4me3, H3K9me3, H3K9ac, H3K9me2, H3K27ac, and H3K27me3. Total histone H3 and IgG occupancy was also analyzed by ChIP-qPCR (**Supplemental Fig S1**).

**Figure 1.**
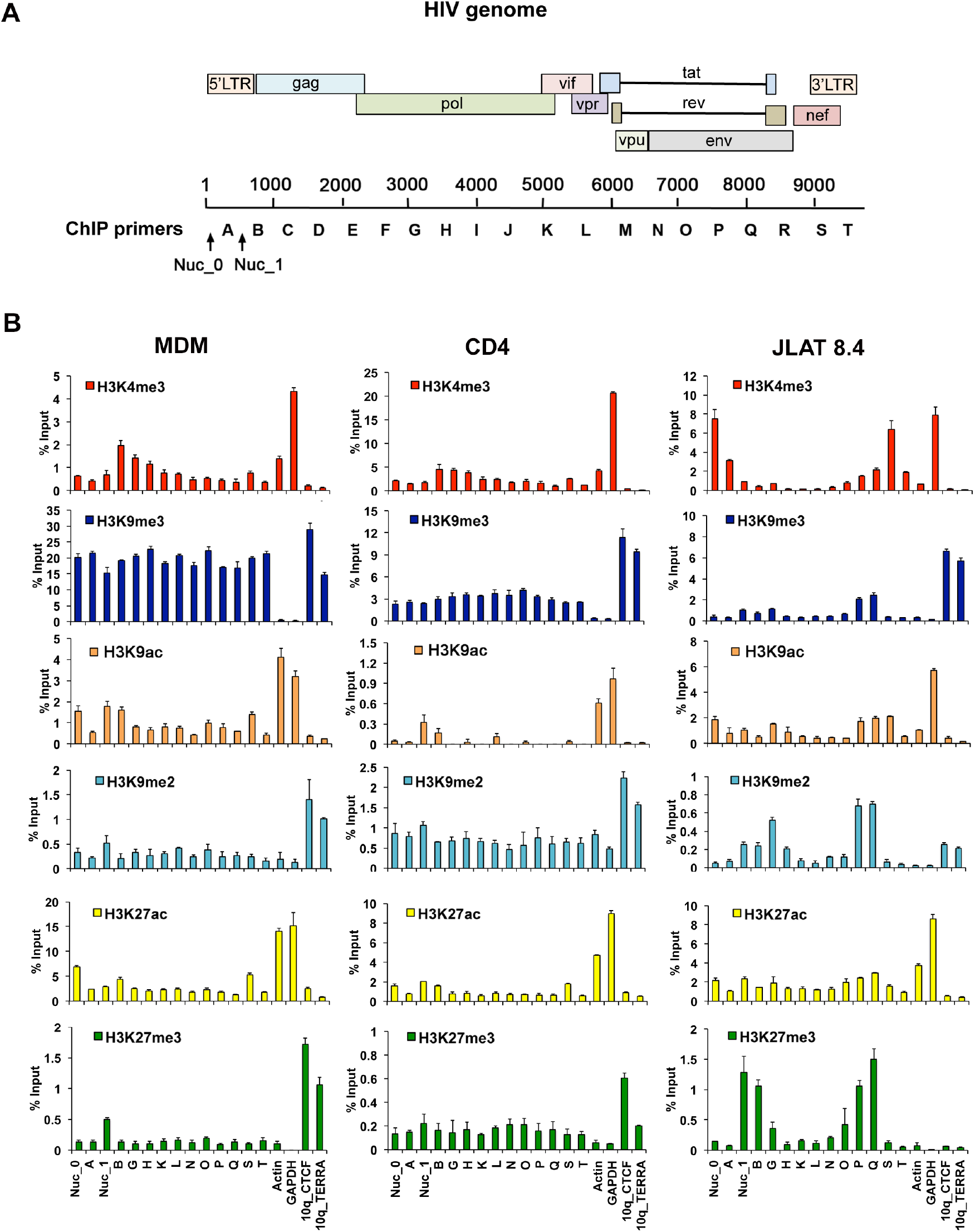
Histone modifications across the HIV genome in MDM, CD4 T cells and JLAT 8.4 cells. **A)** Schematic of HIV genome and positions of primers used for ChIP and DIP. **B)** ChIP-qPCR analysis of the HIV genome in MDM, CD4 T cells, or JLAT 8.4 cells for H3K4me3, H3K9me3, H3K9ac, H3K9me2, H3K27ac, or H3K27me3 using primers spaced across the genome as indicated. Cellular control primers targeted the Actin and GAPDH promoters, and 10q CTCF and 10q TERRA transcript regions. Note that primer stes Nuc0, Nuc1, A, B, S, T target sequences in both 5’ and 3’ LTRs but are depicted graphically with one or the other. Error bars are sdm for 3 technical replicates.

We observed significant differences in the pattern of histone modification between each of these HIV infection models. H3K4me3, a euchromatic mark associated with most transcriptional start sites, was enriched at site B in MDM and T cells, and in JLAT 8.4 at Nuc_0, site A and site S within the nef region and LTR. Nucleosome positions have been well characterized for HIV LTR in JLAT cells (33, 34). H3K9me3, a mark associated with constitutive heterochromatin, was highly enriched across the entire HIV genome in MDM (approaching 20% input), while relatively lower levels were observed in latently infected JLAT 8.4, and intermediate levels in CD4 T cells. H3K27me3, a facultative heterochromatic mark associated with polycomb-mediated repression, was enriched in JLAT 8.4 at Nuc 1 and site B, as well as within the body of the genome within the env gene (P and Q regions). H3K9ac was detected in the Nuc 0 and 1, and site B in MDM, with lower levels in JLAT 8.4 and CD4 T cells. Controls for H3K4me3 and H3K9ac were enriched at active cellular genes Actin and GAPDH as expected, and heterochromatic marks for H3K9me3 or H3K27me3 were enriched at telomeric sites (10q CTCF and TERRA) as expected (with the exception that JLAT 8.4 lacked H3K27me3 enrichment at telomeres). IgG levels were generally below threshold significance at all primer positions with the exception of Nuc 1 in all three cells and positions G, P, Q in JLAT 8.4 which showed a low background signal (**Supplemental Fig S1**). These findings indicate that HIV genomes in MDM are subject to distinct histone modification patterns compared to acute and latent CD4 T cell infection models. We also note that some of these epigenetic marks are enriched in HIV regions outside of the LTR.

### Atypical bivalent histone modification of the MDM LTR

To further explore the ChIP data, we re-analyzed the histone modifications focusing only on the LTR and adjacent nucleosome 2 (site B) for each infection model (**Fig. 2)**. We observed that the LTR in MDM was highly enriched with histone H3K9me3 and to a lesser extent with H3K27ac. The LTR in CD4 T cells also showed a relatively high enrichment of H3K9me3, as well as H3K9me2, and the highest enrichment in H3K4me3 at site B, and the lowest density of total H3 relative to the other cell types tested. JLAT 8.4 cells were highly enriched for H3K4me3 at Nuc_0 and had low levels of H3K9me2 and H3K9me3. H3K27me3 was enriched at Nuc1 and site B in JLAT relative to the other cell types. This latter pattern of histone modifications has been reported previously for JLAT (35–37) and corresponds with bivalent (H3K4me3/K27me3) histone modification observed in some developmentally regulated genes and pluripotent stem cells (38). In contrast to JLAT, the H3K4me3 enrichment in CD4 peaked at site B, located close to nucleosome 2. The relatively high H3K9me3 seen in MDM and CD4 cells, combined with H3K4me3 in CD4 T cells and to a lesser extent in MDM, has been observed at lineage specific regulatory elements, such as methylation pause sties in adipocytes (39). The combination of H3K9me3 and H3K27ac that we find in MDM has also been observed for transposable elements in ES cells (40).

**Figure 2.**
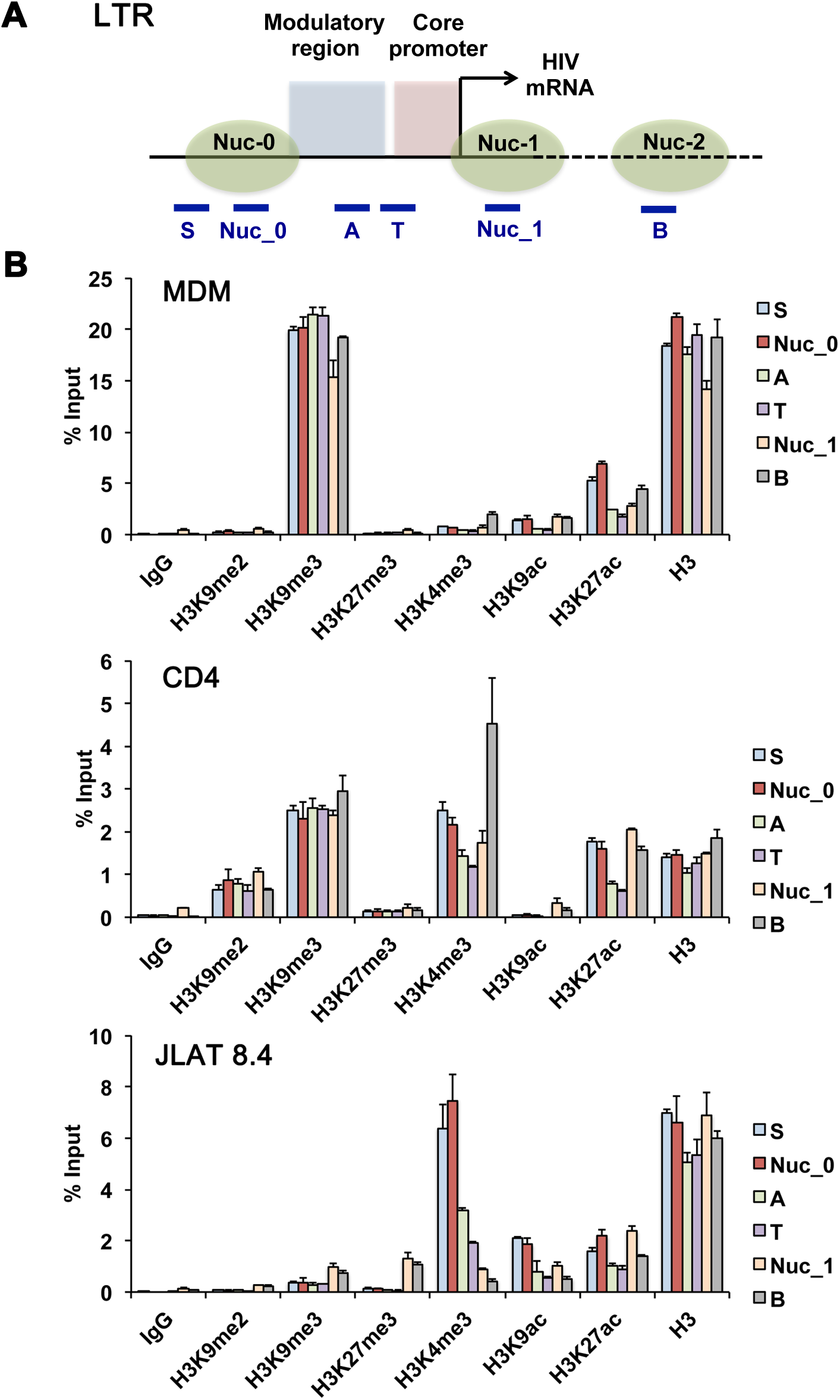
ChIP-qPCR focusing on HIV LTR. **A)** Schematic of HIV 5’ LTR with adjacent nucleosome 2 and positions of primers used in Fig 2B. Solid line represents LTR DNA, and dashed line represents HIV genomic region downstream of 5’ LTR. Primers that amplify sequence duplicated in 5’ and 3’ LTRs are listed. **B)** ChIP-qPCR data from Fig 1 was re-graphed to directly compare enrichments of each histone modification across the LTR region and adjacent nucleosome 2 site for MDM, CD4, and JLAT8.4. Error bars are sdm for 3 technical replicates.

### Antisense transcripts in infected MDM

Antisense transcripts are known to be a source of generating H3K9me3 (41), and antisense transcripts have been reported for HIV particularly during latent infection (42, 43). Therefore, we assayed for anti-sense HIV transcripts using HIV-specific antisense RT primers at nucleotide positions 15 (AS RT-2), 2944 (AS RT-1), and 7431(AS RT-3) in conjunction with qPCR primer pairs positioned at 351-461(primer 2), 3161-3275 (primer 1), 7846-7733 (primer 3) and 8333-8426 (primer 4) based on the YU2 HIV genome (**Fig. 3A**). Antisense transcripts were detected at all positions in YU2-infected MDM and CD4 T cells but not in uninfected control cells (**Fig. 3B**). Similarly, antisense transcripts were not detected in reactions lacking RT (**Fig. 3C**). The relative levels of anti-sense transcript in MDM and CD4 T cells correlated with the relative ratio of sense transcripts for *nef* and *tat* (**Fig. 3D**). These findings indicated that both MDM and CD4 T cells have measurable levels of anti-sense HIV and raise the possibility that antisense transcription contributes to H3K9me3 formation, as has been observed for many endogenous retroviruses (44).

**Figure 3.**
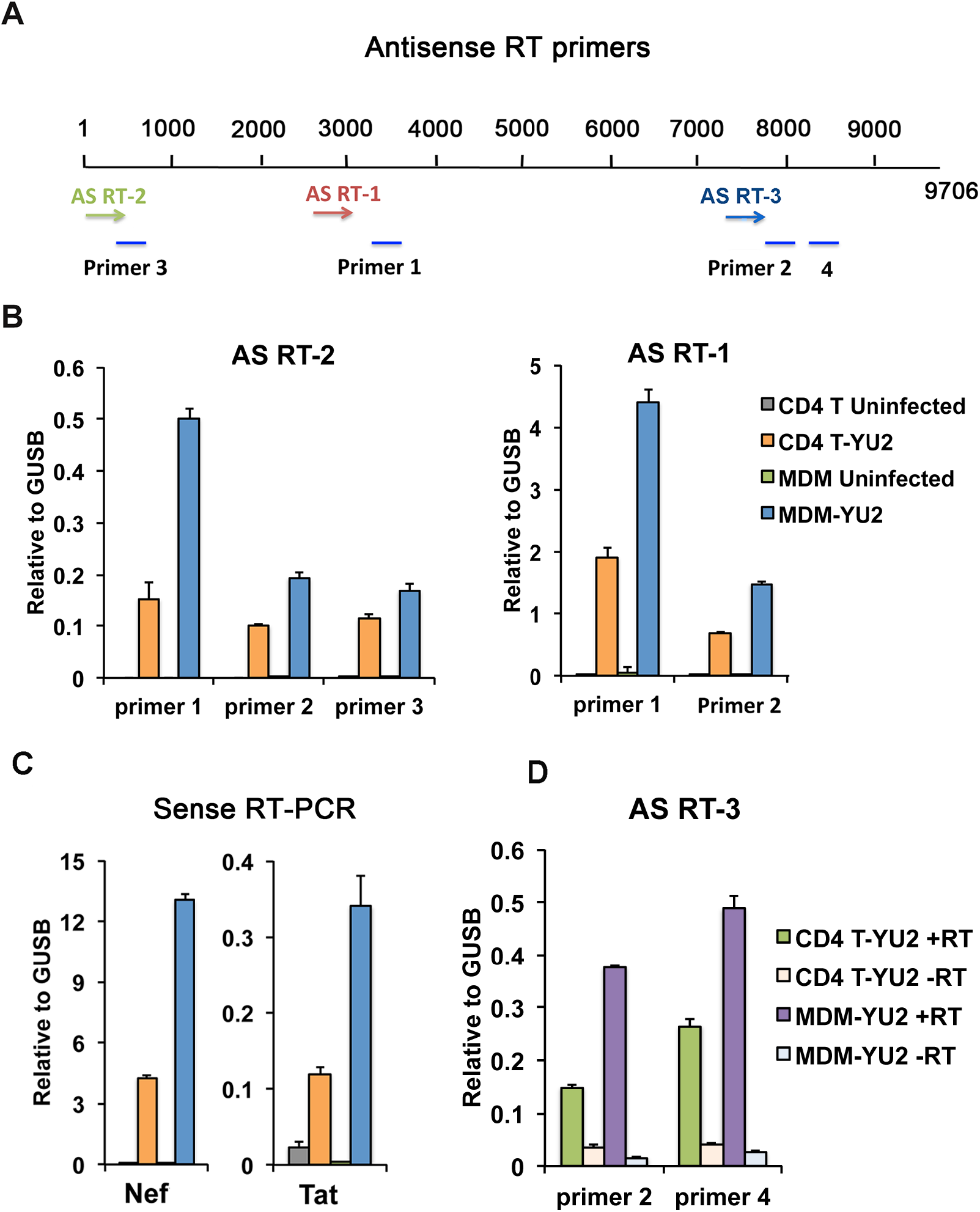
Antisense RNA transcripts of HIV in MDM and CD4 infection. **A)** Schematic of HIV genome showing position of three HIV-specific antisense reverse transcription primers (AS-RT1,2, or 3) and four qPCR primer pairs (1, 2, 3, and 4) used for antisense transcript quantification. **B)** RT-qPCR of antisense transcripts in uninfected and YU2-infected CD4 T cells and MDM, calculated relative to cellular GUSB sense transcript. **C)** RT-qPCR of antisense transcripts in YU2-infected CD4 T cells and MDM with or with addition of RT. **D**) Sense transcription of nef or tat for same infection and RNA samples as shown in panel B.

### Cytosine methylation and hydroxymethylation of HIV genomes

To investigate whether DNA modifications differed in MDM and CD4 T cell infection, we assessed the levels of cytosine methylation (5mC) and hydroxymethylation (5hmc) of HIV DNA in MDM, CD4 T cells, and JLAT 8.4 using MeDIP or hMeDIP assays (**Fig. 4**). We detected elevated levels of 5mC on the JLAT 8.4 genome, including locations at site B, which was previously identified as a CpG island adjacent to the 5’LTR in JLAT (35), and the 3’ regions within the *env* gene (P, Q, R). Relatively low or undetectable levels of 5mc were found associated with HIV infection in MDM and CD4 T cells. In contrast, both MDM and CD4 T cells showed a broad pattern of hydroxymethylation (5hmc) across the HIV genomes, whereas this modification was mostly absent from JLAT 8.4. Cellular controls for 5mC and 5hMC were highly enriched at cellular telomeric positions, and absent in actively transcribed genes Actin and GAPDH. These findings indicate the 5mC is formed in long-term latently infected JLAT cells but is generally not formed during productive infection of primary MDM or CD4 T cells, while 5hmc forms during infection of MDM and CD4 T cells with actively transcribing HIV genomes.

**Figure 4.**
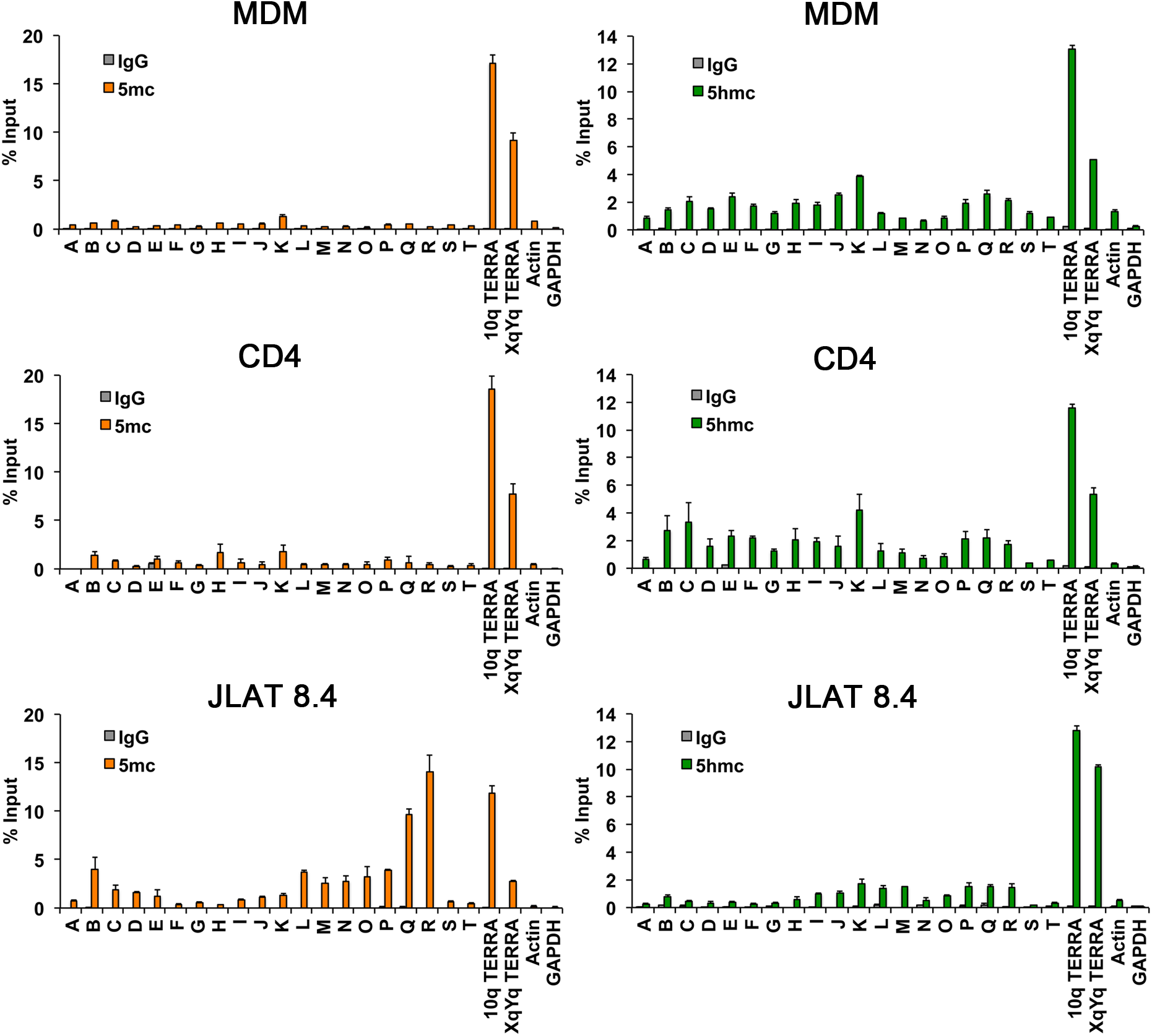
DNA immunoprecipitation (DIP) assay for 5mc or 5hmc across the HIV genome for infected MDM, CD4 T cells, or JLAT 8.4 cells. MeDIP for 5mC (left panels, orange bars) or hmeDIP for 5hmc (right panels, green bars) or control IgG were analyzed using PCR primers across HIV genome, as indicated in Fig. 1A. Cellular gene control sites at 10q or XqYq TERRA or promoter regions for Actin or GAPDH.

### Differential expression of histone and nuclear viral response proteins in MDM and CD4 T cells

To assess whether these epigenetic variations were associated with cell-type specific differences in global histone modifications, we assayed total cellular histone modification levels with or without HIV-1 YU2 infection by Western blot analysis (**Fig. 5A**). We found that histone levels were generally less abundant in MDM relative to CD4 (normalized to Actin and total protein), although these modifications did not change significantly upon HIV infection. Among the histone modifications, H3K9me2 appeared depleted in MDM relative to CD4, and H3K9me3 showed a slight increase (~1.47 fold) upon HIV infection in MDM. We next assayed chromatin proteins that have been implicated in nuclear antiviral functions (**Fig. 5B**). We found that PML had a different distribution of slow-mobility isoforms in HIV infected MDM that were not detectable in CD4 T cells. These slower mobility PML isoforms may reflect PML isoforms and post-translational modifications induced by virus infections in the nucleus (45). Both DAXX were DNMT3A were less abundant in MDM relative to CD4 T cells, but were not affected by HIV infection. In contrast, the methylcytosine oxidase TET2 was downregulated in HIV-infected MDM, but not CD4 T cells. This is consistent with previous studies showing that TET2 is a target of ubiquitin-mediated degradation by HIV Vpr in macrophages (46, 47), and suggests this effect is cell type-specific. We also found that Lamin A/C is expressed at much higher levels in MDM relative to CD4, while the reverse occurs for Lamin B1 expression. These differences may reflect the very different nuclear morphology and cell-cycle properties of MDM and CD4 T cells. HIV also induced p53 in MDM, but not in CD4 T cells (**Fig. 5C**). And strikingly, we found that MDM had near undetectable levels of PARP1, although HIV induced total cellular levels of poly-ADP ribose (PAR), suggesting that other PARPs may be activated in MDM in response to HIV infection. Taken together, these findings underscore substantial differences in nuclear protein biology in MDM and CD4 T cells, and their distinct responses to HIV infection.

**Figure 5.**
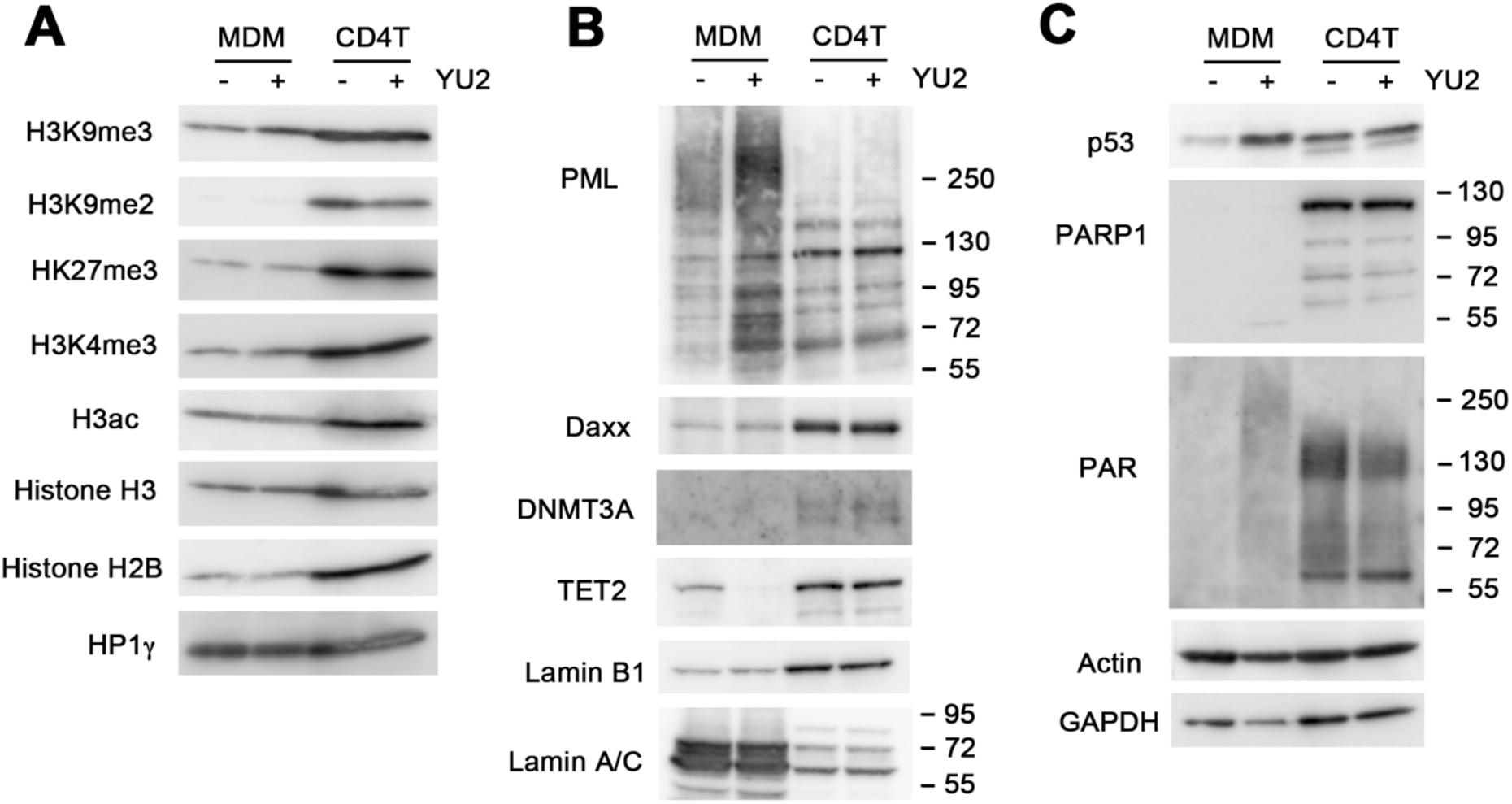
Western blot analysis of MDM and CD4 T cells with HIV-1 YU2 or mock infection. **A**) Westerns were probed for a panel of histone modifications H3K9me3, H3K9me2, H3K27me3, H3K4me3, H3ac, H3, H2B, and HP1γ. **B)** Westerns probed for PML, Daxx, DNMT3A, TET2, Lamin B1, Lamin A/C. **C)** Westerns probed for p53, PARP1, PAR, Actin or GAPDH.

### HIV induced changes in MDM nuclear organization

MDM form multinucleated and giant cells in response to activation signals and in response to HIV infection both *in vitr*o and in the brain *in vi*vo (8, 48–50). We examined the changes in MDM nuclear morphology before and after HIV infection, using immunofluorescence microscopy to detect nuclear proteins and viral p24 Gag antigen (**Fig. 6**). HIV infection led to a marked increase in multinucleated cells enriched with H3K4me3 (~2.0 fold) and 5-hmc (~1.6 fold) (**Fig. 6A, B, and E**). We also observed an increase in punctate PML and DAXX colocalized nuclear bodies in each of the multiple-nuclei in infected MDM (**Fig. 6C, D and E**). We also observed that 5hmc signal changed substantially upon HIV infection in MDM (**Fig. 6F**). In the absence of infection, most 5hmc appeared perinuclear, while after HIV infection 5hmc was strongly enriched in each of the nuclei of the multinucleated MDM. These findings demonstrate that HIV infection remodels MDM nuclear morphology, induces an antiviral response increase in PML nuclear bodies, and induces a strong nuclear relocalization of 5hmc.

**Figure 6.**
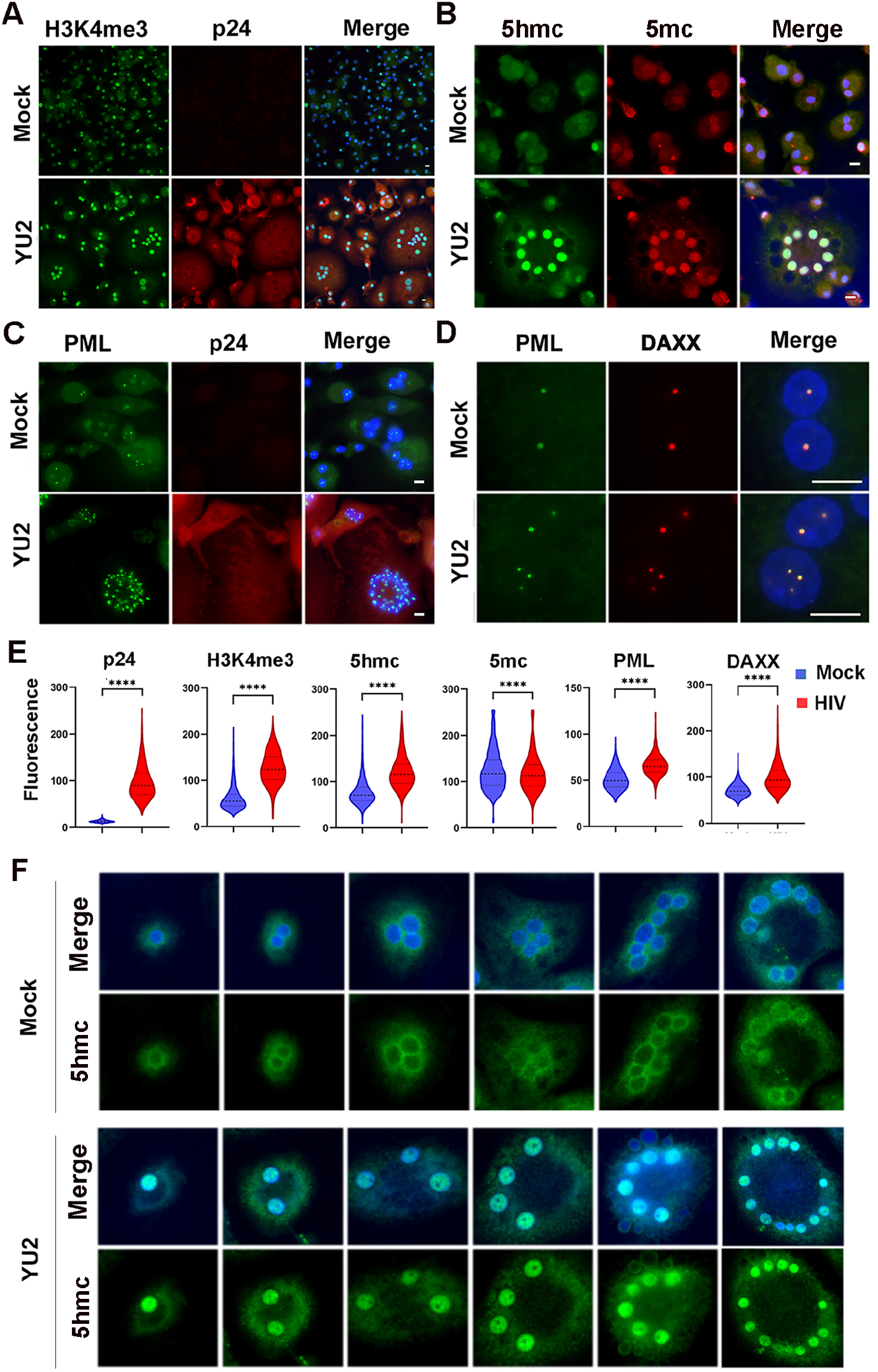
Immunofluorescence microscopy of MDM with HIV-1 YU2 or mock infection. **A)** YU2 or mock infected MDM stained with H3K4me3 (green) or HIV protein p24 (red), merge with Dapi (blue). **B)** 5-hmc (green), 5-mc (red) or merge with Dapi (blue). **C)** PML (green), p24 (red), or merge with Dapi (blue). **D)** PML (green), DAXX (red), merge with Dapi (blue). The images were taken with 20X (**A** – **C**) or 100X (**D**) lens. Scale bar = 10 μm. **E)** Quantification of fluorescence intensity for images represented in panels A-D are provided for cells n>100 with distribution around mean intensity. P values were determined by two-tailed student-t test. ****p<0.0001. **F)** IF for 5hmc in MDM mock (top panels) or YU2 infected (lower panels) with 5hmc (green) or DAPI (blue, merge). Images taken with 20x lens.

## Discussion

While much is known about HIV infection and latency in CD4 T cells, relatively less is known about the regulation of HIV in macrophages. Here, we examined the epigenetic features of HIV in primary human MDMs, employing a brain-derived HIV-1 primary isolate relevant to *in vivo* infection, and compared it with CD4^+^ T cells and the latently infected T cell line JLAT 8.4. Our comparison suggests that MDM use different mechanisms than CD4^+^ T cells to regulate HIV infection. We found HIV sequences in MDM are enriched with H3K9me3 throughout the viral genome, with an atypical bivalent histone modification characterized by high H3K9me3 in combination with H3K27ac (**Fig.7**). We also observed that 5’-hydroxymethylated cytosine (5-hmc) was enriched across the HIV genome in MDM and CD4, but not in JLAT, which were elevated in 5-mc. These data suggest that the epigenetic regulatory features in MDMs are different than those observed in both productively infect CD4 T cells and latently infected JLAT cells, and potentially distinct from previously characterized macrophage activation states.

**Figure 7.**
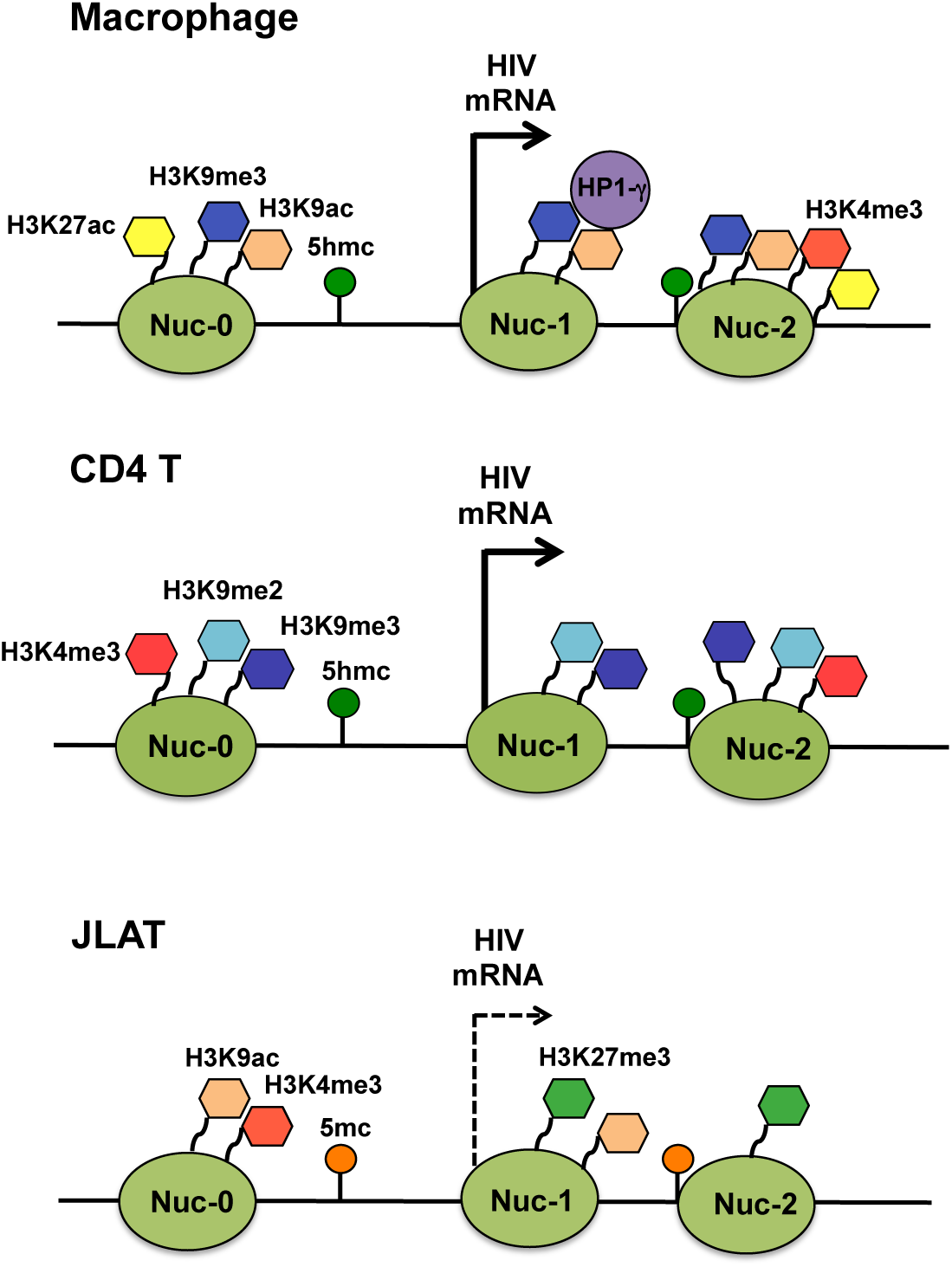
Model of HIV epigenetic regulation. Summary of histone tail modifications associated with LTR region in HIV infected MDM, CD4 T cells and JLAT 8.4 infection.

Our findings are similar to previous reports of H3K9me3-associated heterochromatin that was associated with silencing of HIV LTR post-integration (51, 52). However, in MDM infection where HIV is actively transcribing, the H3K9me3 does not appear to confer transcriptional silencing. In CD4 T cells, H3K9me3 formation was found to depend on the histone methyltransferase SUV39H1 or the SETDB1-associated HUSH complex (53). In microglial cells, transcription factor C/EBP alpha, co-repressor CTIP2, and histone demethylase LSD1 have all been implicated in the formation of silent heterochromatin at the HIV LTR (52, 54, 55). COUP-TF and CTIP2 were found to recruit HP1alpha and H3K9me3 methylation in microglial cells (23, 56, 57). HIC1 and HMGA1 were also found to be chromatin-associated repressors of HIV transcription in microglial cells (58). Whether these same factors function to generate H3K9me3 in MDMs is unknown. Epigenetic control of HIV transcription and replication is also likely to depend on the surrounding chromatin environment. We noted that HIV genomic regions outside the LTR, and especially within the regions encompassing the env ORF, have distinguishing epigenetic marks. In JLAT 8.4, this region was enriched for H3K27me3 and 5-mc. This raises the possibility that regulatory control outside the 5’ LTR may contribute to the epigenetic regulation of HIV.

DNA methylation is also known to contribute to HIV silencing of the 5’ LTR in JLAT and CD4+ T cells (35, 59). We found high levels of 5mC at the LTR in JLAT, as expected, but no significant 5mC in MDM. On the other hand, 5hmc levels were enriched in MDM across the entire HIV genome. To our knowledge, hydroxymethylcytosine has not yet been described in the regulation of HIV in macrophage or T-cells. HIV Vpr has been shown to cause ubiquitin-mediated degradation of TET2 (46, 47), the major enzyme responsible for enzymatically converting methylcytosine to hydroxymethylcytosine in hematopoietic cells (60). Our data show that TET2 is selectively degraded in MDM infection, but not in CD4 T cells (**Fig. 5**). We also observed increased intensity and relocalization of 5hmc from the nuclear periphery to the nucleus in HIV infected MDMs (**Fig. 6**). A similar localization of 5hmc to the nuclear periphery was observed in developing mouse retinal photoreceptor cells (61). How the HIV-dependent degradation of TET2 is balanced with the enrichment and redistribution of 5hmc on the HIV genome remains to be elucidated.

Total levels of histone proteins were lower in MDMs, as were most modified histones, relative to CD4+ T cells. This may reflect the post-mitotic state of MDMs relatively to cycling CD4+ T cells. However, the amount of H3K9me3 relative to other histone modifications appeared to be higher in MDM. While HIV infection did not alter global levels of any histone modification, it did induce many other changes in MDM proteins. In addition to the loss of TET2, HIV induced several modifications associated with viral infection and DNA damage response, including modification of PML, induction of p53, and the generation of PAR. PARP1 has been found to form a complex with Vpr (62), but our findings suggest that PARP1 is undetectable prior to HIV infection in MDM cells. Other PARPs, such as PARP2 or Tankyrase may be responsible for PARylation in MDM cells.

Imaging studies revealed that HIV infection leads to a large increase in multinucleated giant cells, which is well described *in vitro* and *in vivo*. These form in response to macrophage activation due to direct interaction with pathogens (63), or phagocytic substrates (64). For HIV-infected cells, multinucleate giant cells may also result from Env-mediated cell-cell fusion. More recent studies suggest that macrophage multinucleation can arise from mitotic polyploidy and chromothripsis, and not exclusively as a result of cellular fusions (65). Our findings are consistent with the induction of DNA damage based on the increase in phosphorylated p53 and PAR that correlates with the formation of the multinucleated macrophages after HIV infection.

HIV transcriptional regulation has been shown to be controlled primarily by factors that bind to the LTR and the TAR regions, including nucleosomes positioned in close proximity to the transcription start site. Recent studies suggested unintegrated HIV DNA, especially 2-LTR circles were associated with repressive chromatin structure (66, 67), and that circular HIV persists in macrophages since they are non-dividing cells (68). Most of the assays in this study do not distinguish between 2-LTR circles and integrated HIV provirus, so it remains possible that some of the epigenetic signals associated with the HIV genome in MDM are derived from the non-integrated 2-LTR circles. It is also known that most HIV genomes integrate into transcriptionally active loci in the cellular genome, and therefore are not subject to local heterochromatic repression. However, it is not fully known how long-term epigenetic suppression occurs in T-cells, nor how the H3K9me3 repressive chromatin is formed, especially in MDMs where HIV transcription is not silenced. Others have found that H3K9me3 can be associated with transcriptional activation (69, 70) and mRNA elongation (71), especially when this histone modification paired with H3K9ac or H3K4me2 (72). H3K9me3 is also found coupled with H3K27ac at transposable elements that can be transcriptionally activated in some cell types and stress conditions (40). We provide evidence that HIV genomes have H3K9me3 distributed throughout the viral genome, and that antisense transcription occurs that may initiate at the 3’ LTR. The detection of antisense transcription in MDM is consistent with the model that antisense transcripts can recruit factors that generate H3K9me3 heterochromatin (35, 36). Paradoxically, the elevated H3K9me3 in MDM does not correspond to transcriptional repression, suggesting that H3K9me3 may not serve as transcriptionally repressive heterochromatin in the context of the HIV genome in macrophage. It is likely that additional repressive factors that restrict RNA polymerase II function may be not fully operational at the HIV LTR in MDM infection.

## Materials and Methods

### Cells and viral infection

Monocytes were purified from healthy donors and cultured in Iscove’s Modified Dulbecco’s Medium (IMDM) containing 10% human AB serum with penicillin, streptomycin and 1% glutamine. Monocytes were maintained in culture for 7 days to allow differentiation into monocyte-derived macrophage (MDM) prior to HIV-1 infection. CD4 T cells were purified from healthy donors and cultured in RPMI1640 containing 10% FCS with penicillin, streptomycin and 1% glutamine. T cells were stimulated with 5 μg/ml phytohemagglutinin (PHA) for 3 days prior to HIV-1 infection, then treated with 10 ng/ml interleukin-2 (IL-2). Cell purification used negative selection employing the Rosette-Sep platform (Stemcell Technologies) and were carried out by the Penn CFAR Immunology Core. All data represent a minimum of three independent experiments using cells from different donors.

The brain-derived HIV-1 pYU2 infectious molecular clone (IMC) with all accessory genes intact (31) was provided by Dr. B. Hahn (University of Pennsylvania), and infectious virus generated by transfection of 293T cells. Virus was treated with DNase (to prevent inadvertent transfection of residual plasmid during infections) and quantified by HIV-1 p24 Gag antigen by ELISA. To enhance entry into CD4 T cells, the HIV-1 YU2 IMC was co-transfected with plasmid encoding VSVg to generate mixed pseudotypes, harvested and quantified similarly. To enhance infection of MDM in the Chromatin and DNA Immunoprecipitation (ChIP, DIP) experiments, transduction with the Vpx protein of SIV was carried out. Vpx-containing pseudotype virions were generated by co-transfecting 293T cells with plasmids encoding SIV Gag, SIV Vpx gene and VSVg (73), provided by J. Skowronski (Case Western Reserve University). Viral particles were harvested 3 days later, quantified by SIV Gag p27 antigen by ELISA, and 3 ng of p27 was used for transduction per 10^6^ MDM at same day of HIV-1 infection. MDM at 7 days post-plating or CD4 T cells 3 days post-PHA stimulation were infected with HIV-1 YU2 using 7 ng of viral p24 Gag antigen per 10^6^ MDM or 3.5 ng of viral p24 Gag antigen per 10^6^ CD4 T cells. Cells were infected by spinoculation at 1200 g for 2 hours at room temperature, and then maintained in culture until analyzed.

For ChIP, MeDIP and hMeDIP assays, 50 nM of the RT inhibitor efavirenz (EFV) was added to MDM 6 days post infection to restrict further rounds of re-infection and enable maximal integration, or added to CD4 T cells 4 days post infection, then cells were cultured for additional 4 days before harvest. For RT-PCR, EFV was added to MDM cells 7 days post infection, cells were then cultured for additional 7 days before harvest. EFV was added to CD4 T cells 2 days post infection, then cells were cultured for additional 2 days before harvest. For Western blot analysis, cells were harvested 7 days post infection.

### Chromatin immunoprecipitation (ChIP)-qPCR assays

ChIP-qPCR assays were performed as described previously (74). Quantification of precipitated DNA was determined using real-time PCR and the delta Ct method for relative quantitation (ABI 7900HT Fast Real-Time PCR System). Rabbit IgG (2729S, Cell Signaling), anti-H3K4me3 (07-473, Millipore Sigma), H3K9me2 (C15410060, Diagenode), H3K9me3 (C15410056, Diagenode), H3K9ac (07-352, Millipore Sigma), H3K27me3 (C15410069, Diagenode), H3K27ac (ab4729, Abcam), and pan-histone H3 (07-690, Millipore Sigma) were used in ChIP assays. Primers for ChIP and DIP assays are listed in Supplemental Table S1.

### MeDIP and hMeDIP assays

Total genomic DNA was purified using Wizard® Genomic DNA Purification Kit (Promega, A1120) then subjected to methylcytosine-DNA-immunoprecipitiation (MeDIP) or hydroxymethylcytosine-DNA-IP (hMeDIP) assays. The MeDIP or hMeDIP assays were performed using MagMeDIP kit (Diagenode, C02010021) or hMeDIP kit (Diagenode, C02010031). Quantification of precipitated DNA was determined using real-time PCR and the delta Ct method for relative quantitation (ABI 7900HT Fast Real-Time PCR System).

### Western blot assay

Rabbit anti-H3K4me3, H3K9me2, H3K9me3, H3K27me3 and pan histone H3 antibodies used in Western blotting were same as antibodies used in ChIP assays. Rabbit polyclonal anti-H3ac (06-599, Millipore Sigma), anti-H2B (07-371, Millipore Sigma), anti-Lamin B1 (12586S, Cell Signaling), anti-TET2 (21207-1-AP, Proteintech), anti-PARP-1 (210-302-R100, Alexis), anti-Daxx (D7810, Millipore Sigma), anti-GAPDH (Cell signaling 2118); Mouse monoclonal anti-p53 (OP43, Millipore Sigma), anti-PML (ab96051, Abcam), anti-Lamin A/C (MANLAC1, DSHB), anti-PAR (4335-MC-100, Trevigen), anti-HP1γ (MAB3450, Chemicon), anti-DNMT3A (IMG-268A, Imgenex); Actin-Peroxidase antibody (Sigma A3854).

### Immunofluorescence (IF)

On day 8 post HIV infection, cells were washed with 1XPBS and fixed for 15 min with 2% paraformaldehyde (Electron Microscopy Sciences) in 1XPBS, then washed twice with 1XPBS, recovered with 70% ethanol, washed with 1XPBS and permeabilized for 15 min with 0.3% TritonX-100 (Sigma) in PBS. Cells were then incubated in blocking solution (0.2% fish gelatin, 0.5% BSA in 1XPBS) for 30 min, at room temperature (RT). Primary antibodies were diluted in blocking solution and applied on the cells for 1h at RT followed with 1xPBS washing. For 5hmc and 5mc staining, after the permeabilization and PBS-washing steps the cells were treated with 4N HCl for 30 min at room temperature. Cells were then washed with 1xPBS three times, incubated with 5-hmc or 5mc antibodies in blocking solution. Cells were further incubated with fluorescence-conjugated secondary antibodies in blocking solution for 1h, RT, counterstained with Dapi and mounted in Fluoromount-G medium (SouthernBiotech). Images were taken at Nikon Upright Microscope using 20X or 100X lens and processed with Adobe Photoshop CS6. Antibodies used in IF: mouse anti-p24 (ab9071, Abcam), rabbit anti-H3K4me3 (07-433, Millipore Sigma), rabbit anti-PML (A301167A, Bethyl), mouse anti-5mc (C15200081-100, Diagenode), rabbit anti-5hmc (39769, Active motif), goat anti-DAXX (sc-167A, Santa Cruz), AlexaFluor594 or AlexaFluor488 (Invitrogen). Original fluorescence images were captured with standardized acquisition parameters using a Nikon 80i upright microscope with ImagePro Plus software (Media Cybernetics). Fluorescence image intensity was quantified by gathering each set of images into a single multipoint, multichannel ND file using Nikon Elements AR software (Nikon Instruments) and analyzed together. For each individual image, the DAPI channel was used to define the size, shape and location for each nucleus, the outline was converted to a region of interest and applied to all available channels for that image. Mean intensity was then collected in each channel for each region of interest.

### RNA extraction and quantitative RT-PCR

RNA was isolated from 2 x 10^6^ cells using RNeasy plus mini Kit (Qiagen). Reverse transcription was performed with either random decamers or HIV antisense-specific primers. HIV antisense qPCR was carried out using 4 specific primer pairs and the antisense reverse transcription cDNA, while cellular GUSB and HIV tat and nef qPCR was done using random decamer cDNA. Real-time PCR was performed with SYBR green probe in an ABI Prism 7900 and the delta Ct method for relative quantitation. Primers for HIV antisense-specific reverse transcription and RT-qPCR are listed in Supplement Table S2.

## Acknowledgments

This work was supported by NIH grant R61/33 133696 (RGC, PML). We thank J. Skowronski and B. Hahn for constructs. We acknowledge assistance from Andreas Wiedmer and the Wistar Cancer Center Core Facilities in Imaging and Flow Cytometry, and the Immunology and Virology Cores of the Penn Center for AIDS Research (P30-AI045008).

## Supplemental Figure Legends

**Supplemental Figure S1.**
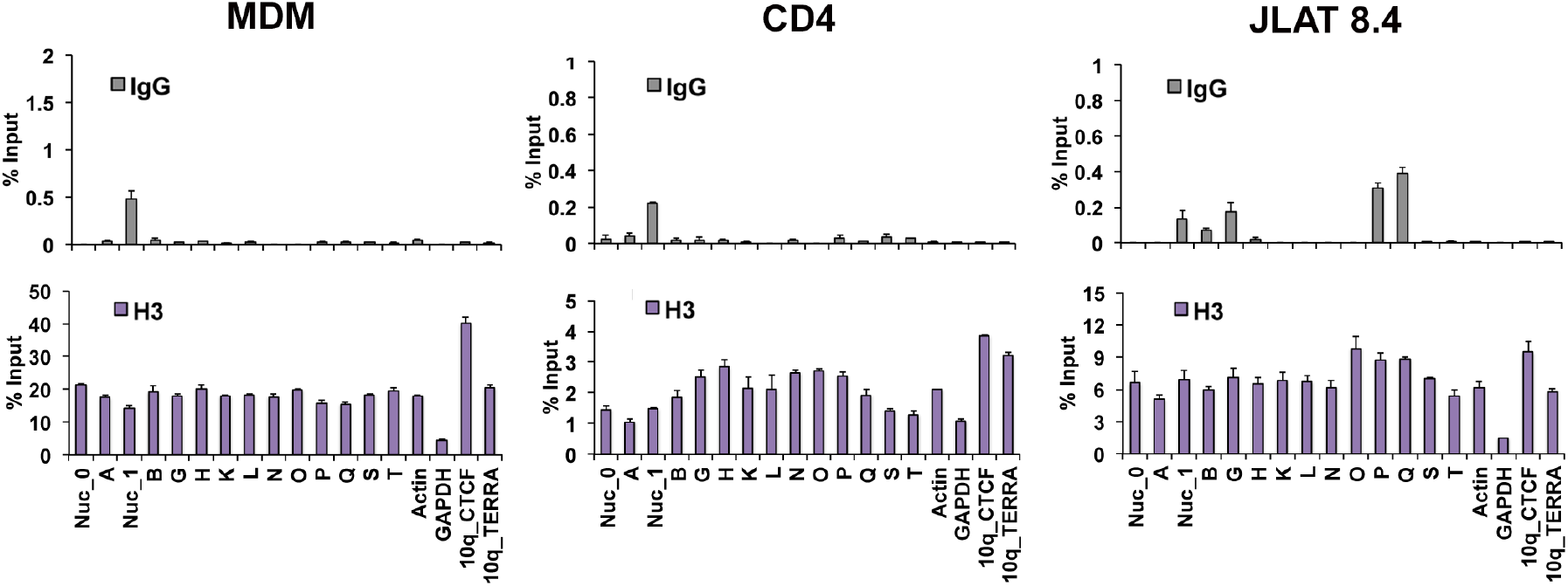
ChiP-qPCR controls. ChIP-qPCR of HIV genome (as in Figure 1) showing control IgG and total H3 antibody in MDM, CD4, and JLAT 8.4.

**Supplemental Table S1.**
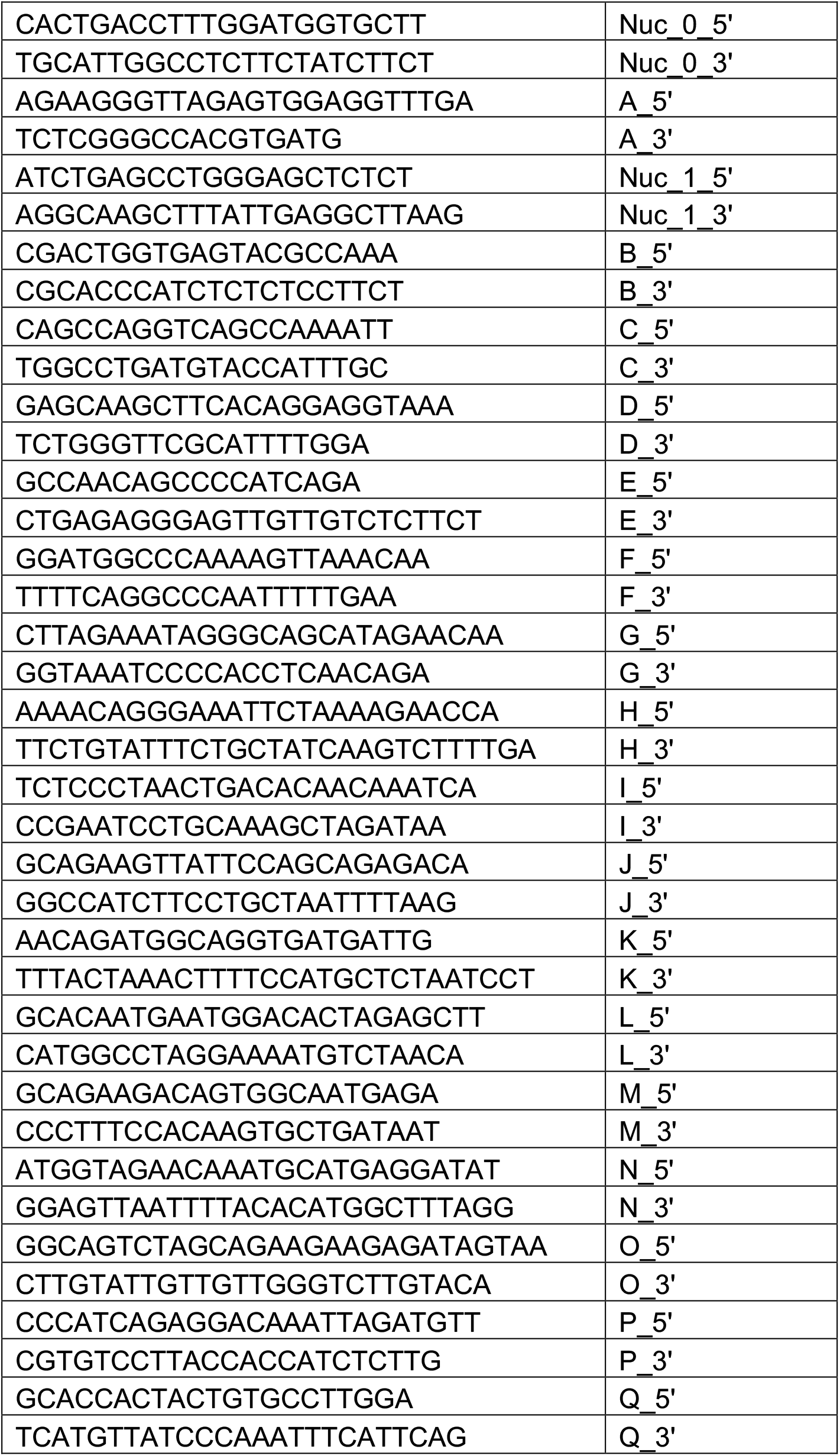
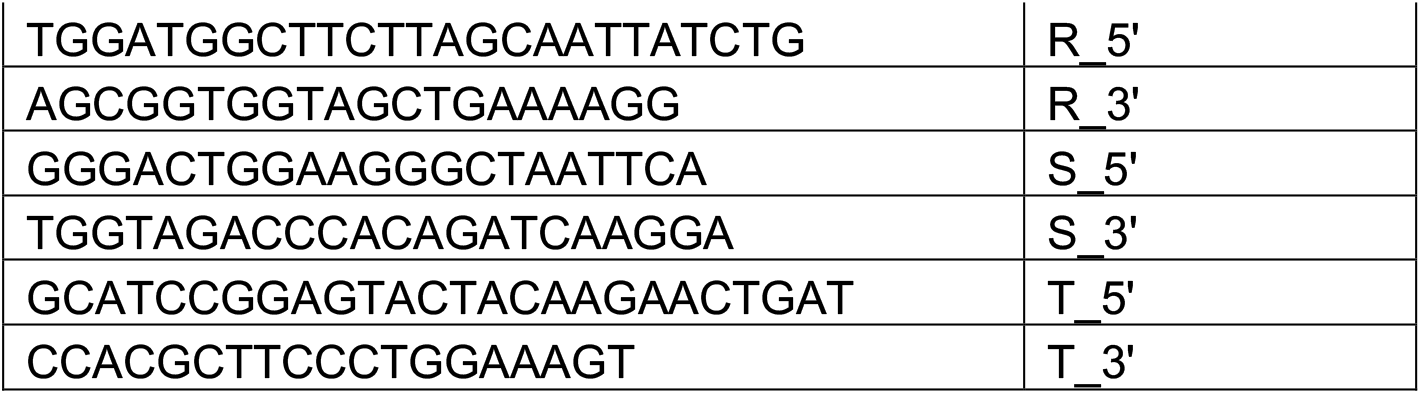
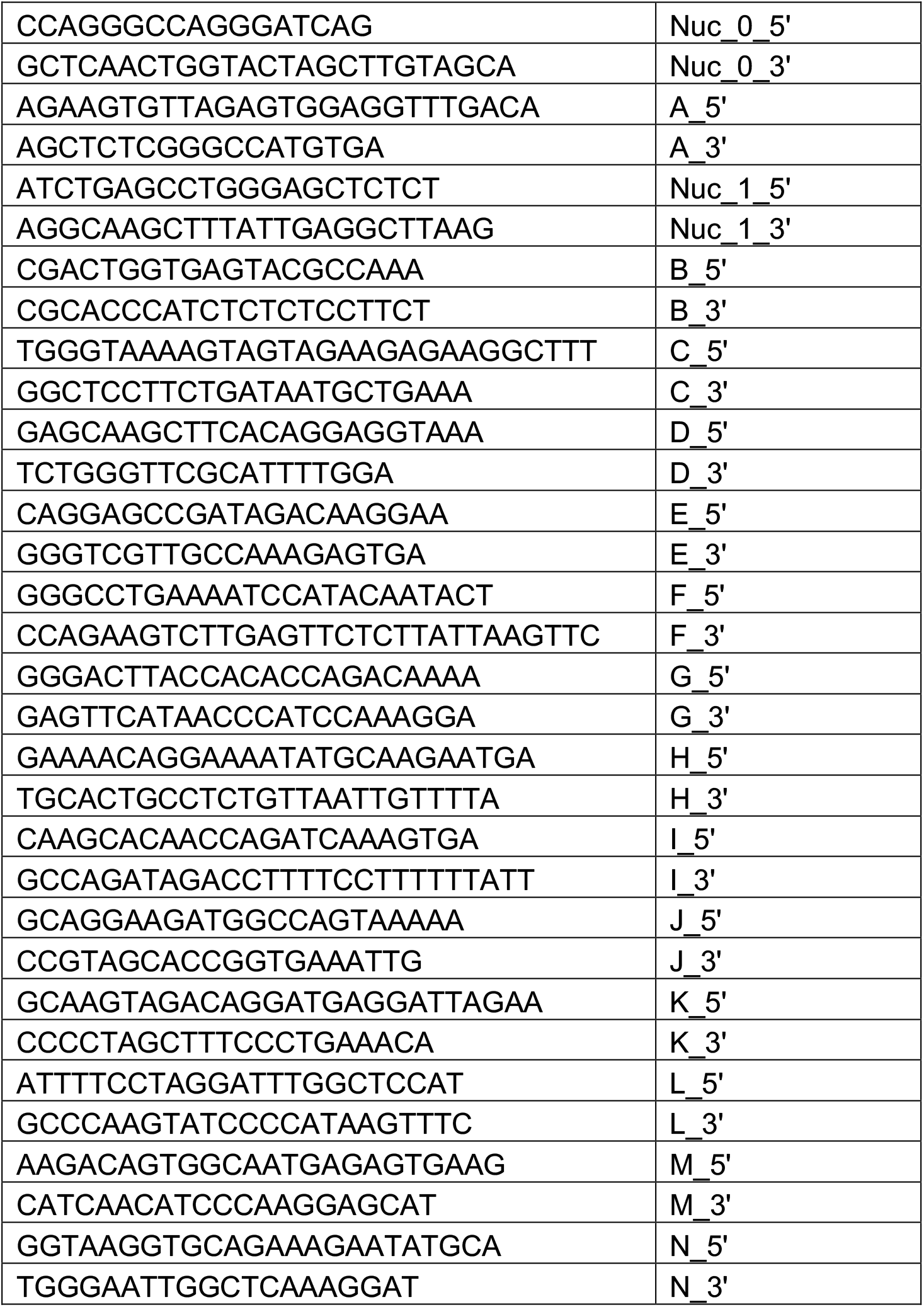
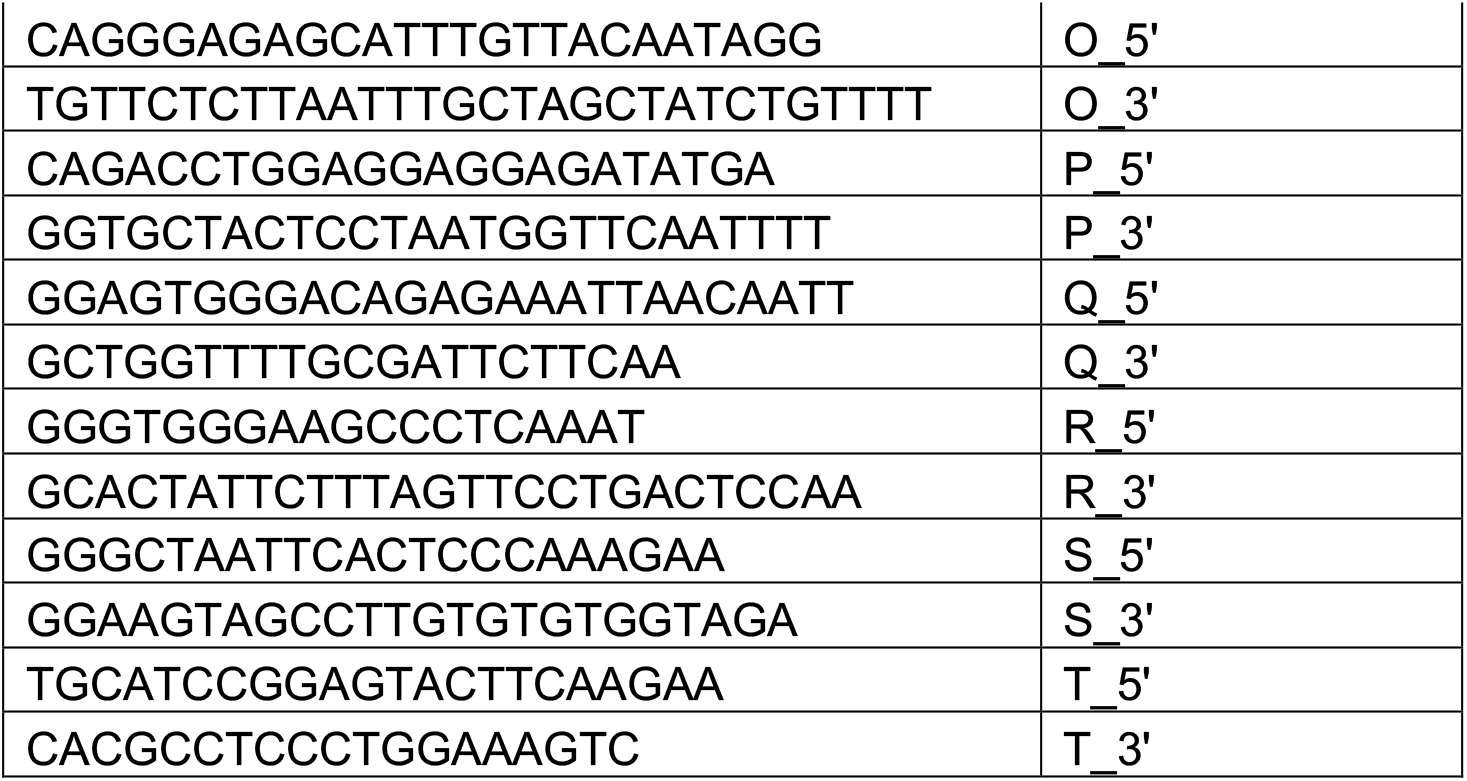
HIV genomic primers. A. Genomic primers for HIV YU2 strain. B. Genomic primers for HIV HXB2 strain (JLAT 8.4).

**Supplemental Table S2.**
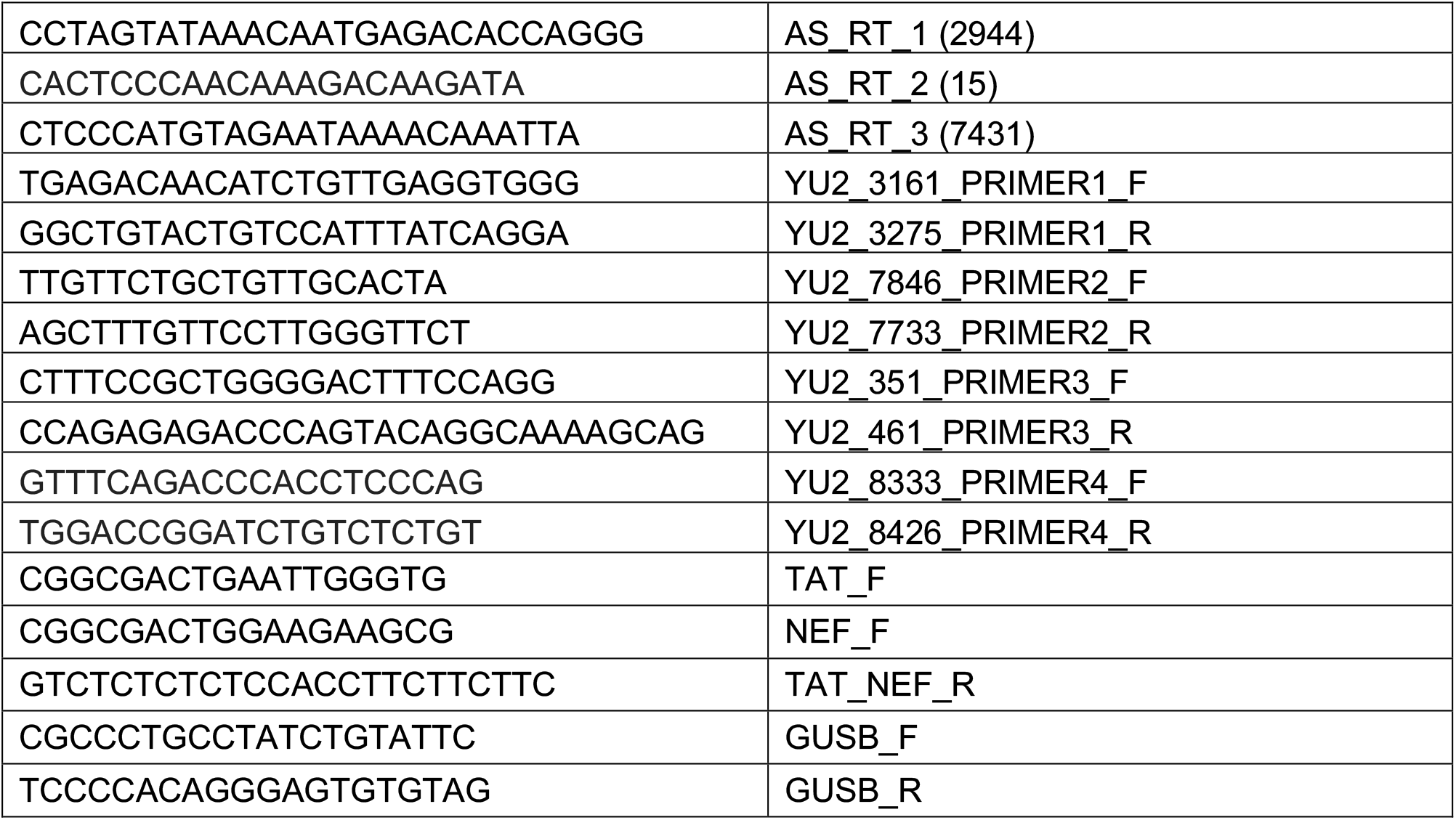
RT-qPCR primers.

